# Pyruvate partitioning defines an immunosuppressive metabolic niche in colorectal cancer

**DOI:** 10.64898/2026.10.16.759773

**Authors:** Jiawei Zheng, Bin Lian, Shiyin Huang, Jinyuan Liu, Yingxin Wu, Huiling Zhai, Yanlin Qiu

## Abstract

**Purpose:** Colorectal cancer (CRC) reprograms glucose metabolism, diverting pyruvate from mitochondrial oxidation toward lactate export. How this partitioning decision is organized across cell types and tissue space, and how it relates to immune programmes, remains unresolved.

**Methods:** We defined a Pyruvate Partitioning Index (PPI) as the difference between an oxidative arm (MPC1, MPC2, PDHA1, PDHB, DLAT, DLD, PDHX) and an export arm (LDHA, LDHB, SLC16A1, SLC16A3, SLC16A7), and computed it across TCGA COADREAD (n = 592), GSE39582 (n = 579), two single-cell atlases (370,115 and 63,689 cells), 13 Visium sections and an exploratory urothelial immunotherapy cohort (IMvigor210, n = 298).

**Results:** PPI was lowest in microsatellite-unstable (MSI) tumours (Mann–Whitney p = 3.8×10^−9^) and inversely associated with overall survival (log-rank p = 1.2×10^−4^). Epithelial cells were the most oxidative compartment (median +0.058) and myeloid cells the most export-prone (−0.171; Kruskal p ≈ 0), reproduced independently; MMRd epithelium was intrinsically export-shifted (−0.012 vs +0.076 vs +0.192 for MMRd, MMRp and normal). Spatially, PPI was non-random (Moran’s I significant in 12/13 sections), with low-PPI spots enriched for SPP1+ macrophages, myofibroblasts and plasma cells, but not CD8+ T-cell factors. Low-PPI tumours showed coordinated increases in immune-checkpoint, myeloid and Treg programmes (checkpoint rho = −0.262), and network perturbation identified LDHA as the hub.

**Conclusions:** Pyruvate partitioning is cellularly and spatially organized in CRC and defines a myeloid-stromal immunosuppressive niche. Because MSI tumours—the best CRC immunotherapy responders—had the lowest PPI, the PPI–response relationship appears cancer-type specific and requires validation in CRC cohorts.

## 1. Introduction

Colorectal cancer is a leading cause of cancer death worldwide, and its outcome remains heterogeneous even within molecularly similar tumours [1]. A hallmark of CRC is metabolic reprogramming, in which glucose-derived pyruvate is diverted from mitochondrial oxidation toward lactate production and export [2,3].

Pyruvate sits at a branch point. It can enter the mitochondrion for oxidative decarboxylation through the pyruvate dehydrogenase (PDH) complex, or be reduced to lactate by LDH and exported through monocarboxylate transporters (MCT1/SLC16A1, MCT4/SLC16A3) [4,5]. The partition between these arms is set by structural genes and determines the extracellular lactate load, which modulates innate and adaptive immunity in the tumour microenvironment [6,7].

Lactate metabolism has emerged as a determinant of the immunosuppressive CRC microenvironment. Prior studies have scored lactate metabolism as an overall activity—for example the four-gene LMscore [8] or a pan-cancer lactate score [9]—or treated single genes such as *MPC1* as prognostic markers [10], and have linked glycolysis to immune evasion in MSI-H CRC [11] and to hypoxia-related subtypes [12]. These analyses, however, quantify lactate metabolism as abundance rather than as the decision between oxidation and export, and none resolve the partition at single-cell and spatial resolution.

This gap matters because the two arms have opposite immunological consequences: oxidation sustains mitochondrial respiration, whereas export generates the lactate that conditions myeloid and stromal cells toward immunosuppression. Whether a single index of pyruvate partitioning can delineate an immune niche in CRC is unresolved.

Here we define a Pyruvate Partitioning Index (PPI) and resolve it across bulk, single-cell and spatial transcriptomics to ask (i) whether PPI is cellularly and spatially organized in CRC, and (ii) whether low PPI marks an immunosuppressive niche.

## 2. Methods

### 2.1 Data sources

Bulk RNA-seq and clinical annotations for TCGA COADREAD (n = 592) were retrieved via cBioPortal; external validation used GSE39582 (n = 579). Single-cell data comprised the Pelka et al. atlas (370,115 cells) [13] and the SMC cohort GSE132465 (63,689 cells, 23 patients) [14]. Spatial transcriptomics used 13 Visium sections from a CRC consensus-molecular-subtype study with pre-computed cell2location factors [15]. The exploratory immunotherapy analysis used IMvigor210 (atezolizumab-treated metastatic urothelial carcinoma, n = 298 with response annotation) [16].

### 2.2 Pyruvate Partitioning Index

PPI was defined as the difference between the oxidative arm (*MPC1, MPC2, PDHA1, PDHB, DLAT, DLD, PDHX*) and the export arm (*LDHA, LDHB, SLC16A1, SLC16A3, SLC16A7*). Each gene was z-scored across samples and arm scores were the mean of member-gene z-scores. In single-cell data, z-scores used global parameters across all cells so that cross-compartment comparison was not artificially zero-centred.

### 2.3 Immune programmes

Immune-checkpoint (PDCD1, CTLA4, LAG3, HAVCR2, TIGIT), cytotoxicity (CD8A, IFNG, GZMB, PRF1, NKG7), myeloid (CD163, CD86) and Treg (FOXP3, IL2RA) programmes were scored as the mean z-score of member genes. These gene sets are disjoint from the PPI panel, so PPI–immune associations are not confounded by shared genes.

### 2.4 Survival, subtyping and association

Overall survival was compared between PPI strata by Kaplan–Meier and log-rank test; univariate and stage-adjusted Cox models were fitted. Associations with molecular subtype and with immune programmes were tested by rank-based correlation (Spearman). Effect sizes are reported as rho throughout, without causal interpretation.

### 2.5 Single-cell and spatial analysis

Cell-type PPI was summarized by median and compared by Kruskal–Wallis; MMRd versus MMRp epithelial PPI by Mann–Whitney. Spatial autocorrelation used Moran’s I with 99 permutations; PPI was correlated with cell2location factors per section and meta-analysed across sections.

### 2.6 Cell–cell communication and network perturbation

Ligand–receptor communication between low-PPI epithelium and myeloid cells was scored with a CellPhoneDB-style product of ligand and receptor expression with 500 permutations of the epithelial PPI labels (FDR < 0.1). For network perturbation, epithelial cells were split into low- and high-expression groups per target gene, a weighted co-expression network (|rho| > 0.15) was built per group, and a degree-difference ratio was computed against 200 random relabels.

### 2.7 Exploratory immunotherapy analysis

As an exploratory analysis, PPI was computed in IMvigor210 from the same 12-gene panel. Response was examined both as a binary variable (CR/PR vs SD/PD; Wilcoxon test) and as the four-level ordinal variable (CR/PR/SD/PD; Kruskal–Wallis test); objective response rate was compared between PPI-stratified groups; overall survival used Kaplan–Meier and Cox models. Association with tumour mutational burden used Spearman correlation. Because IMvigor210 is a urothelial carcinoma cohort, these results are reported as exploratory and are not generalized to CRC.

### 2.8 Software

Analyses used Python 3 and R 4.6 (Scanpy, Squidpy, survival, Biobase). Permutation p-values were floored at permutation resolution. Core analysis scripts are provided in the Supplementary Code (absolute paths replaced by placeholders, input-data provenance documented); the complete analysis pipeline is available from the corresponding author on request.

## 3. Results

### 3.1 PPI stratifies CRC by subtype and outcome

Across TCGA COADREAD (n = 592), PPI was lowest in MSI-H tumours (median −0.454 vs +0.066 for MSS, n = 89 vs 466; Mann–Whitney p = 3.8×10^−9^) and inversely associated with overall survival (log-rank p = 1.2×10^−4^; Cox HR = 0.62, p = 0.002; Fig. 1a,b). A 21-feature ElasticNet Cox model reached a test C-index of 0.66. We note at the outset that MSI tumours— the CRC subgroup with the best immunotherapy response—had the lowest PPI; the implications are addressed in the Discussion.

**Fig. 1.**
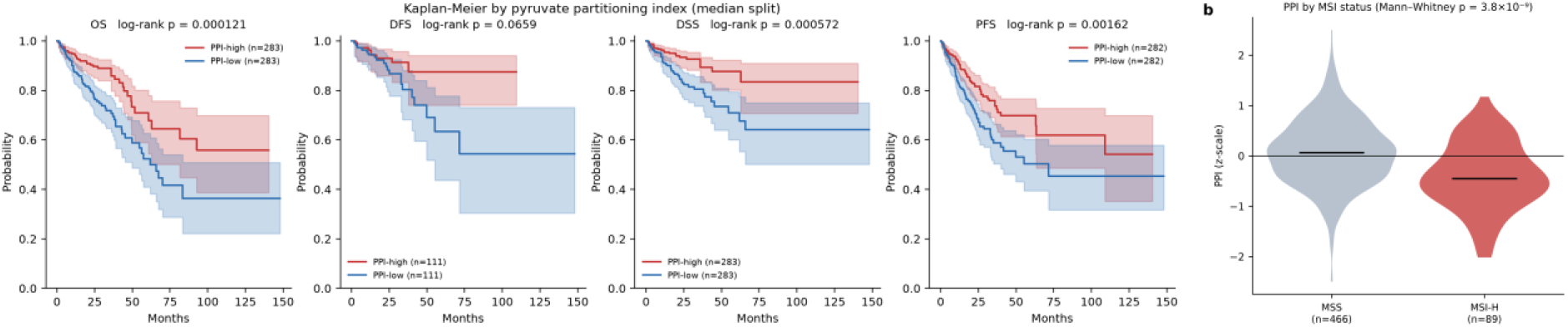
PPI stratifies CRC by outcome and MSI status. (a) Kaplan–Meier curves of TCGA COADREAD patients stratified by PPI for overall, disease-specific, disease-free and progression-free survival (OS log-rank p = 1.2×10^−4^; DSS, DFS and PFS directionally consistent). (b) PPI in MSI-H versus MSS tumours (median −0.454 vs +0.066; n = 89 vs 466; Mann– Whitney p = 3.8×10^−9^).

### 3.2 Pyruvate partitioning is cell-type specific

At single-cell resolution, epithelial cells were the most oxidative compartment (median PPI +0.058), whereas myeloid cells were the most export-prone (−0.171), with T/NK (−0.065), mast (−0.067) and stromal (−0.050) cells intermediate (Kruskal H = 21,060, p ≈ 0; Fig. 2a,b). This ordering was reproduced in the independent GSE132465 cohort (epithelial +0.149, myeloid −0.144).

**Fig. 2.**
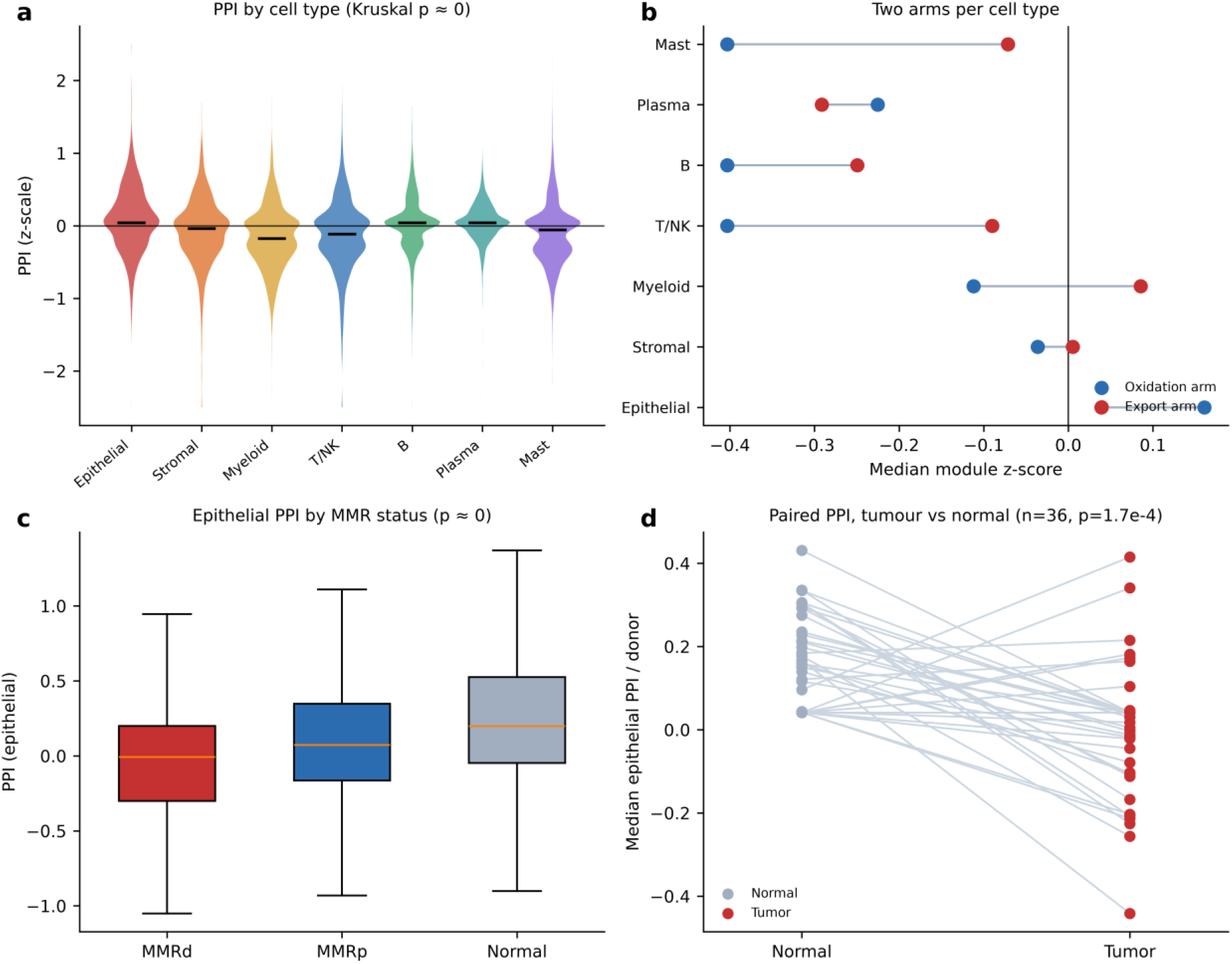
Cell-type and MMR-associated organisation of pyruvate partitioning. (a) Per-cell PPI across the seven major compartments of the Pelka CRC atlas (Kruskal p ≈ 0). (b) Oxidation-arm and export-arm scores per compartment. (c) Epithelial PPI by MMR status. (d) Paired epithelial PPI, tumour versus matched normal (36 donors; Wilcoxon p = 1.7×10^−4^).

### 3.3 MMRd epithelial cells are intrinsically export-shifted

Within epithelium, MMRd tumour cells were export-shifted relative to MMRp and normal cells (−0.012 vs +0.076 vs +0.192, p ≈ 0; Fig. 2c). In paired analysis, malignant epithelium drifted toward export relative to matched normal epithelium (36 donors, +0.024 vs +0.163, Wilcoxon p = 1.7×10^−4^; Fig. 2d), indicating a cell-intrinsic metabolic shift rather than a stromal-composition artefact.

### 3.4 PPI defines a myeloid-stromal spatial niche

PPI was non-randomly organized in space (Moran’s I significant in 12/13 sections; Fig. 3a). Low-PPI spots were reproducibly enriched for SPP1+ macrophages (mean rho −0.098), myofibroblasts (−0.103), IgG+ plasma cells (−0.100) and Stromal 3 (−0.105), each consistent across 12/13 sections, whereas CD8+ T-cell factors showed no spatial association with PPI (mean rho ≈ −0.006; Fig. 3b). Low-PPI regions therefore mark a myeloid-stromal rather than a T-cell-excluded niche.

**Fig. 3.**
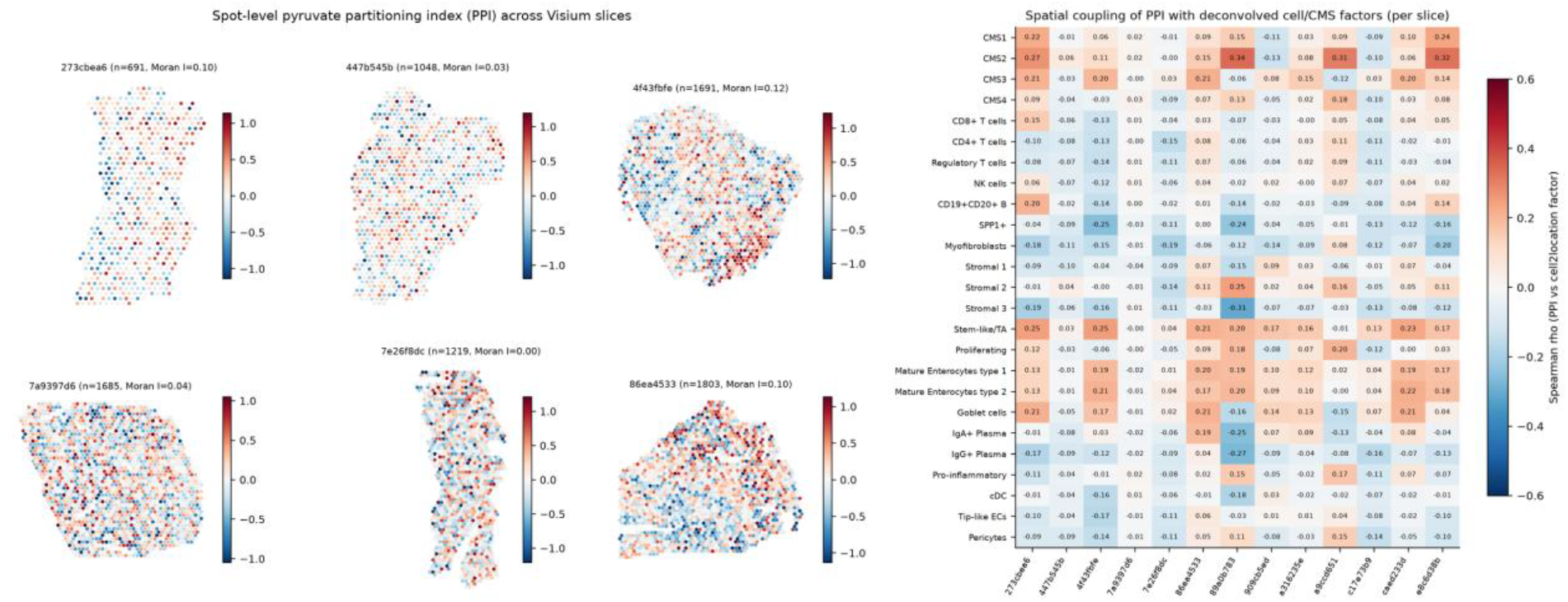
Spatial organisation of PPI. (a) Spot-level PPI across representative Visium sections (Moran’s I with 99 permutations). (b) Per-section Spearman correlation between PPI and cell2location factors (SPP1+ macrophages, myofibroblasts, IgG+ plasma cells, CD8+ T cells).

### 3.5 Patient-level PPI is inversely associated with myeloid infiltration

In GSE132465, patient-level tumour PPI correlated inversely with myeloid fraction (rho = −0.635, p = 0.001, n = 23; Fig. 4a), confirming at the cohort level that low-PPI tumours are myeloid-enriched. This estimate rests on 23 patients and is reported without confidence intervals.

**Fig. 4.**
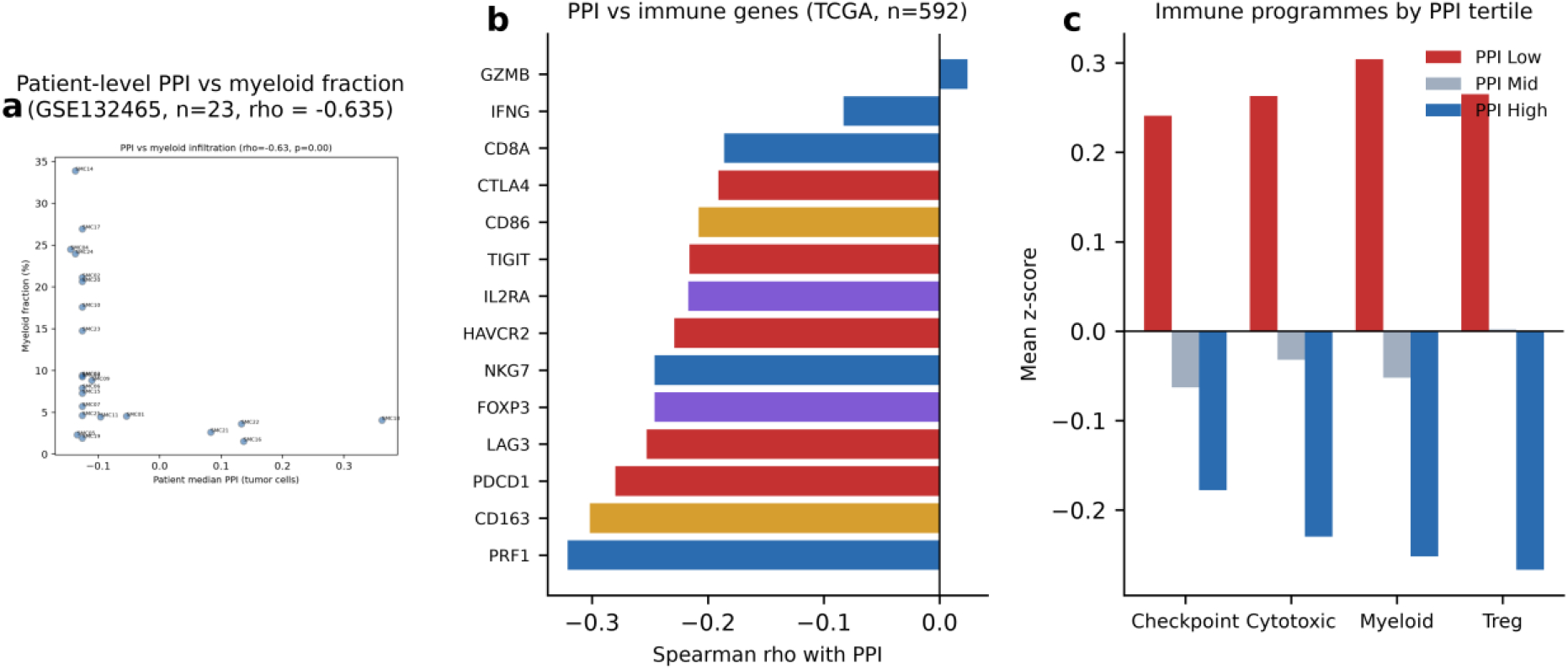
PPI and immune programmes. (a) Patient-level correlation between tumour PPI and myeloid fraction in GSE132465 (n = 23; rho = −0.635). (b) Spearman correlation between PPI and individual immune genes in TCGA (n = 592). (c) Mean immune-programme scores by PPI tertile (checkpoint, cytotoxicity, myeloid, Treg).

### 3.6 Low PPI shows modest coupling to immune-checkpoint and myeloid programmes

Across TCGA (n = 592), PPI correlated negatively with the immune-checkpoint score (rho = −0.262, p = 1.0×10^−10^), the myeloid score (rho = −0.261, p = 1.1×10^−10^), the cytotoxicity score (rho = −0.207, p = 3.9×10^−7^) and the Treg score (rho ≈ −0.25; Fig. 4b). These are weak-to-moderate associations (|rho| ≈ 0.2–0.26): low-PPI tumours show a modest, coordinated shift toward checkpoint, myeloid and Treg programmes, consistent in direction—though not in magnitude—with the myeloid-stromal niche observed spatially.

### 3.7 Exploratory analysis of immunotherapy response in a urothelial cohort

In the independent urothelial carcinoma cohort IMvigor210 (atezolizumab, n = 298), responders had higher PPI than non-responders (median +0.174 vs −0.086; Wilcoxon p = 0.037; Fig. 6a), and the objective response rate was numerically higher in the high-PPI group (0.282 vs 0.174). However, the four-level response comparison (CR/PR/SD/PD) was not significant (Kruskal– Wallis p = 0.223), and overall survival did not differ between PPI strata (log-rank p = 0.46; Cox HR = 0.93, p = 0.66). PPI correlated weakly with tumour mutational burden (rho = +0.195, p = 0.003). Taken together, the association between PPI and response is borderline and detectable only after dichotomizing response; because this cohort is urothelial carcinoma, the finding cannot be generalized to CRC, and it does not override the fact that MSI CRC—the best immunotherapy responders—had the lowest PPI (see Discussion).

### 3.8 Low-PPI epithelium emits immunosuppressive and chemotactic signals

Cell–cell communication analysis identified 31 of 71 ligand–receptor axes as significantly stronger from low-PPI epithelium (Fig. 5a), dominated by alarmin and chemotactic signals (ANXA1→FPR1; S100A9→TLR4), angiogenesis (VEGFA→NRP1/FLT1),immunosuppression (TGFB1→TGFBR1/2) and myeloid recruitment (CXCL3→CXCR2; CCL20→CCR6). SPP1 was only marginally upregulated in low-PPI epithelium (log2FC +0.06), indicating that SPP1+ macrophages arise primarily from recruited myeloid cells rather than from epithelial secretion [21].

**Fig. 5.**
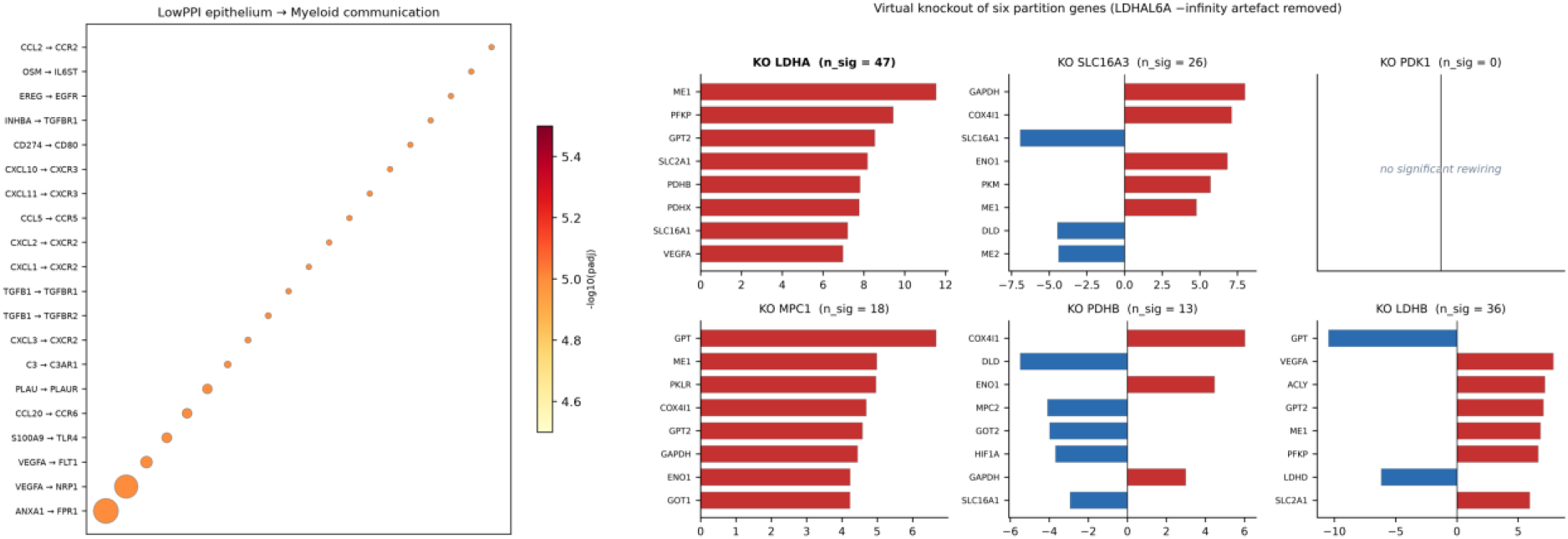
Communication and network perturbation. (a) Ligand–receptor axes significantly stronger from low-PPI epithelium (31 of 71 axes; FDR < 0.1). (b) Degree-difference ratio after network perturbation of six partition genes (*LDHA, SLC16A3, PDK1, MPC1, PDHB, LDHB*); *LDHA* produced the largest rewiring, whereas *PDK1* produced none.

**Fig. 6.**
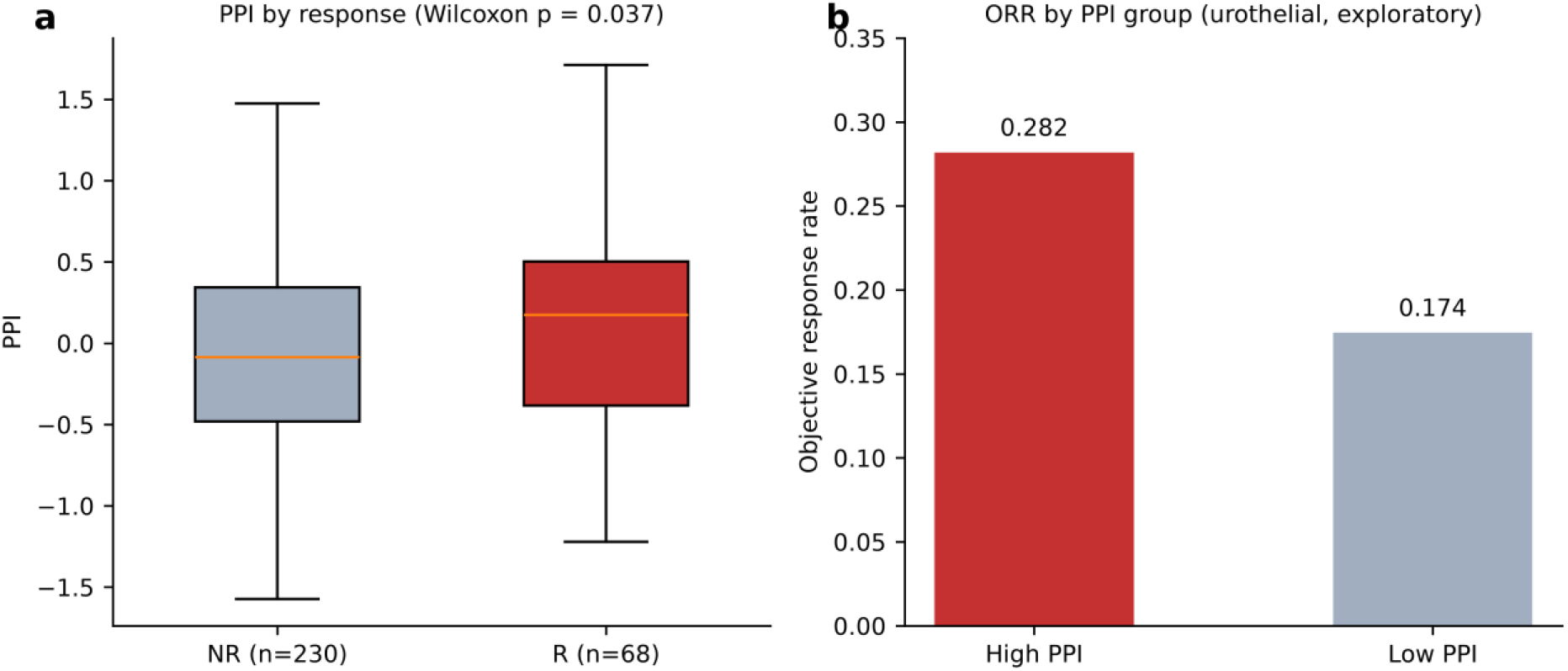
Exploratory analysis of PPI and immunotherapy response (urothelial carcinoma, IMvigor210). (a) PPI in responders versus non-responders (Wilcoxon p = 0.037). (b) Objective response rate by PPI group. Note: exploratory, non-CRC cohort; not generalizable to CRC.

### 3.9 LDHA is the hub of the partition

Network perturbation identified *LDHA* as the dominant hub, with its high-expression state rewiring 47 genes—including the malic enzyme *ME1* (z = +11.5), *PFKP* and *GPT2*—linking glycolysis, anaplerosis and oxidation (Fig. 5b). *PDK1* produced no significant rewiring, indicating that partition topology is set by structural genes rather than the PDK1 regulator.

## 4. Discussion

These findings indicate that pyruvate partitioning is cellularly and spatially organized in CRC, where it defines a myeloid-stromal niche associated with modest enrichment of checkpoint, myeloid and Treg programmes—consistent with recent single-cell and spatial mapping of metabolically defined malignant states in CRC [30]. The claim is observational and correlational.

The myeloid-biased export phenotype is consistent with reports that tumour-associated macrophages take up lactate and undergo lactate-driven reprogramming [17], and the association of low PPI with poorer survival agrees with lactate-metabolism scoring studies [8,12]. Unlike the four-gene LMscore, which quantifies overall lactate-metabolism activity [8], PPI quantifies the balance between oxidation and export; and unlike single-gene *MPC1* analyses [10], it places the gene inside a two-arm difference. The correspondence between metabolic and immune classifications in CRC has been noted previously [28].

Mechanistically, the coupling of low PPI to checkpoint/myeloid/Treg programmes is consistent with a lactate-driven immunosuppressive circuit. Lactate inhibits effector T-cell function [6], promotes M2-like macrophage polarization [18] and drives protein lactylation that reinforces immunosuppressive transcription [19,20]. In CRC specifically, SPP1+ macrophages—the population enriched in low-PPI niches here—drive T-cell dysfunction and immunotherapy resistance through an SPP1-SOCS1-IFN axis [21], are linked to peritoneal immunotherapy resistance [22] and to metastasis [23], and analogous metabolically defined immune niches have been described in other tumour types [29]. Targeting lactate export (MCT4) augments checkpoint blockade in CRC models [24], and MCT4 expression itself has prognostic and immunological significance across cancers [25].

An important tension must be stated plainly. MSI tumours had the lowest PPI (Results 3.1), yet MSI-H/dMMR CRC is the tumour type with the best established response to immune checkpoint blockade. A low-PPI ⇒ poor-response model would therefore make the wrong prediction for MSI CRC. Several factors may resolve this. First, bulk PPI is a mixed signal: because myeloid and lymphoid cells are intrinsically export-prone (Results 3.2), immune-infiltrated MSI tumours will have a lowered bulk PPI partly for compositional reasons rather than through tumour-intrinsic metabolism alone. Second, in CRC, immunotherapy benefit is driven primarily by antigenicity (mismatch repair status, mutational and neoantigen burden) [2,3] rather than by metabolic state, so a metabolic index would not be expected to dominate. Third, our only response data come from a urothelial cohort, where the PPI–response association (borderline, p = 0.037 on dichotomized response) may not transfer across cancer types. We therefore do not claim that PPI stratifies immunotherapy response in CRC; the exploratory analysis is reported to motivate—not to substitute for—dedicated validation in CRC immunotherapy cohorts, and it should be interpreted alongside established response biomarkers such as immune signatures and myeloid composition [26,27].

The principal advantage of this framework is its multi-scale consistency within CRC: the cell-type axis, the MMRd cell-intrinsic shift, the spatial myeloid-stromal niche and the patient-level myeloid association are each supported by independent datasets. The exploratory immunotherapy analysis is presented as hypothesis-generating only.

Several limitations apply. The associations are correlational; PPI is sensitive to panel composition and standardisation; bulk PPI mixes tumour and immune compartments; the spatial autocorrelation, while significant, is modest (Moran’s I up to 0.18); the patient-level myeloid correlation rests on 23 patients without confidence intervals; immune-programme associations are weak in magnitude (|rho| ≈ 0.2–0.26); and the immunotherapy analysis was performed in a urothelial carcinoma cohort and is not generalizable to CRC. No wet-lab validation has been performed.

Future work should (i) deconvolve bulk PPI into tumour-intrinsic and microenvironmental components, (ii) test whether export-arm inhibition (LDHA/MCT4) remodels the myeloid-stromal niche in CRC models, and (iii) evaluate PPI in dedicated CRC immunotherapy cohorts, with MSI status as a stratification variable.

## Statements and Declarations

### Ethics approval and consent to participate

This study analysed publicly available, de-identified datasets. It does not contain clinical studies or patient data requiring ethical approval or informed consent.

### Consent for publication

Not applicable.

### Availability of data and materials

All data analysed are publicly available: TCGA COADREAD via cBioPortal; GSE39582 and GSE132465 via GEO; the Pelka single-cell atlas and the Visium sections via CZ CELLxGENE; IMvigor210 via the IMvigor210CoreBiologies resource. Processed result files summarising every headline number in this manuscript are provided as Electronic Supplementary Material.

### Code availability

Core analysis scripts are provided as Electronic Supplementary Material (absolute paths replaced by placeholders; input-data provenance documented in the accompanying README). The complete analysis pipeline is available from the corresponding author on request.

### Author contributions

Conceptualization: Jiawei Zheng, Bin Lian. Formal analysis: Jiawei Zheng, Shiyin Huang. Methodology: Jinyuan Liu, Jiawei Zheng. Software: Jiawei Zheng, Jinyuan Liu. Validation: Shiyin Huang, Huiling Zhai. Investigation: Yingxin Wu, Yanlin Qiu.

Data curation: Yingxin Wu, Shiyin Huang. Visualization: Jiawei Zheng, Huiling Zhai. Resources: Yanlin Qiu. Supervision: Bin Lian. Project administration: Bin Lian. Writing – original draft: Jiawei Zheng. Writing – review and editing: all authors. All authors read and approved the final manuscript.

### Use of AI-assisted tools

Large language models were used for language editing and to assemble reference metadata (author lists, volume/issue/page numbers), which was subsequently checked against PubMed records. All analyses, data interpretation and conclusions were performed and verified by the authors.

## References

1. Guinney J, Dienstmann R, Wang X, de Reyniès A, Schlicker A, Soneson C, et al. The consensus molecular subtypes of colorectal cancer. Nat Med. 2015;21(11):1350–6.

2. Becht E, de Reyniès A, Giraldo NA, Pilati C, Buttard B, Lacroix L, et al. Immune and Stromal Classification of Colorectal Cancer Is Associated with Molecular Subtypes and Relevant for Precision Immunotherapy. Clin Cancer Res. 2016;22(16):4057–66.

3. Cohen R, Heran M, Pudlarz T, Hilmi M, Tournigand C, André T, et al. Going beyond microsatellite instability for immunotherapy in metastatic colorectal cancer: Consensus molecular subtypes and tumor mutational burden. Bull Cancer. 2019;106(2):151–161.

4. McFate T, Mohyeldin A, Lu H, Thakar J, Henriques J, Halim ND, et al. Pyruvate dehydrogenase complex activity controls metabolic and malignant phenotype in cancer cells. J Biol Chem. 2008;283(33):22700–8.

5. Woolbright BL, Rajendran G, Harris RA, Taylor JA 3rd. Metabolic Flexibility in Cancer: Targeting the Pyruvate Dehydrogenase Kinase:Pyruvate Dehydrogenase Axis. Mol Cancer Ther. 2019;18(10):1673–1681.

6. Wang ZH, Peng WB, Zhang P, Yang XP, Zhou Q. Lactate in the tumour microenvironment: From immune modulation to therapy. EBioMedicine. 2021;73:103627.

7. Gao Y, Zhou H, Liu G, Wu J, Yuan Y, Shang A. Tumor Microenvironment: Lactic Acid Promotes Tumor Development. J Immunol Res. 2022;2022:3119375.

8. Lan H, Li Y, Li H, Guo J, Wang C, Tang Q. Single-cell and spatial transcriptomics reveal lactate-active epithelial-immune cell crosstalk that reprograms the immune microenvironment in colorectal cancer. Transl Oncol. 2026;71:102864.

9. Yin TT, Huang MX, Wang F, Jiang YH, Long J, Li L, et al. Lactate score predicts survival, immune cell infiltration and response to immunotherapy in breast cancer. Front Genet. 2022;13:943849.

10. Zhang E, Zhou X, Fan X, Li S, Ding C, Hong H, et al. Exploring the relationship between lactate metabolism and immunological function in colorectal cancer through genes identification and analysis. Front Cell Dev Biol. 2023;11:1173803.

11. Li C, Cai P, Zeng H, Li J, Hu H, Zhang J, et al. Glycolysis-Driven Immune Evasion in Microsatellite Instability-High Colorectal Cancer: An Integrated Single-Cell and Spatial Transcriptomics Study. Onco Targets Ther. 2025;18:1027–1042.

12. Huang A, Sun Z, Hong H, Yang Y, Chen J, Gao Z, et al. Novel hypoxia- and lactate metabolism-related molecular subtyping and prognostic signature for colorectal cancer. J Transl Med. 2024;22(1):587.

13. Pelka K, Hofree M, Chen JH, Sarkizova S, Pirl JD, Jorgji V, et al. Spatially organized multicellular immune hubs in human colorectal cancer. Cell. 2021;184(18):4734–4752.

14. Lee HO, Hong Y, Etlioglu HE, Cho YB, Pomella V, Van den Bosch B, et al. Lineage-dependent gene expression programs influence the immune landscape of colorectal cancer. Nat Genet. 2020;52(6):594–603.

15. Valdeolivas A, Amberg B, Giroud N, Richardson M, Gálvez EJC, Badillo S, et al. Profiling the heterogeneity of colorectal cancer consensus molecular subtypes using spatial transcriptomics. NPJ Precis Oncol. 2024;8(1):10.

16. Mariathasan S, Turley SJ, Nickles D, Castiglioni A, Yuen K, Wang Y, et al. TGFβ attenuates tumour response to PD-L1 blockade by contributing to exclusion of T cells. Nature. 2018;554(7693):544–548.

17. Yan C, Yang Z, Chen P, Yeh Y, Sun C, Xie T, et al. GPR65 sensing tumor-derived lactate induces HMGB1 release from TAM via the cAMP/PKA/CREB pathway to promote glioma progression. J Exp Clin Cancer Res. 2024;43(1):105.

18. Wang L, Ma Y, Zhang S, Yang Y, Huang B. NFATc2 promotes lactate and M2 macrophage polarization through USP17 in lung adenocarcinoma. Anticancer Drugs. 2024;35(5):385–396.

19. Zhang D, Tang Z, Huang H, Zhou G, Cui C, Weng Y, et al. Metabolic regulation of gene expression by histone lactylation. Nature. 2019;574(7779):575–580.

20. Xie B, Zhang M, Li J, Cui J, Zhang P, Liu F, et al. KAT8-catalyzed lactylation promotes eEF1A2-mediated protein synthesis and colorectal carcinogenesis. Proc Natl Acad Sci U S A. 2024;121(8):e2314128121.

21. Sun L, Chu X, Kong T, Chen X, Ru J, Gao Q, et al. An SPP1-SOCS1 pathway constrains interferon responses in tumor-associated macrophages and shapes an immunosuppressive tumor microenvironment. Immunity. 2026;59(5):1422–1437.e9.

22. Dai X, Lai C, Hong L, Jin Y, Cheng J, Yan H, et al. Evolution of SPP1(+) cavity macrophage-mediated immunotherapy resistance in peritoneal metastasis of colorectal cancer. Cell Rep Med. 2026;7(8):102930.

23. Liu X, Qin J, Nie J, Gao R, Hu S, Sun H, et al. ANGPTL2+cancer-associated fibroblasts and SPP1+macrophages are metastasis accelerators of colorectal cancer. Front Immunol. 2023;14:1185208.

24. Babl N, Decking SM, Voll F, Althammer M, Sala-Hojman A, Ferretti R, et al. MCT4 blockade increases the efficacy of immune checkpoint blockade. J Immunother Cancer. 2023;11(10).

25. Du W, Zang B, Wo Y, Chen S. SLC16A3 (MCT4) expression in tumor immunity and Metabolism: Insights from pan-cancer analysis. Biochem Biophys Rep. 2025;42:102034.

26. Ferreiro-Pantín M, Anido-Herranz U, Betancor YZ, Cebey-López V, León-Mateos L, García-González J, et al. Clinical, molecular, and immune correlates of the Immunotherapy Response Score in patients with advanced urothelial carcinoma under atezolizumab monotherapy: analysis of the phase II IMvigor210 trial. ESMO Open. 2023;8(4):101611.

27. Zeng D, Ye Z, Wu J, Zhou R, Fan X, Wang G, et al. Macrophage correlates with immunophenotype and predicts anti-PD-L1 response of urothelial cancer. Theranostics. 2020;10(15):7002–7014.

28. Soldevilla B, Carretero-Puche C, Gomez-Lopez G, Al-Shahrour F, Riesco MC, Gil-Calderon B, et al. The correlation between immune subtypes and consensus molecular subtypes in colorectal cancer identifies novel tumour microenvironment profiles, with prognostic and therapeutic implications. Eur J Cancer. 2019;123:118–129.

29. Chu T, Zhu G, Tang Z, Qu W, Yang R, Pan H, et al. Metabolism archetype cancer cells induce protumor TREM2(+) macrophages via oxLDL-mediated metabolic interplay in hepatocellular carcinoma. Nat Commun. 2025;16(1):6770.

30. Gui K, Yang T, Xiong C, Wang Y, He Z, Li W, et al. STC2+ Malignant Cell State Associated with EMT, Tumor Microenvironment Remodeling, and Poor Prognosis Revealed by Single-Cell and Spatial Transcriptomics in Colorectal Cancer. Oncol Res. 2025;34(1):24.

